# Stabilization of spine Synaptopodin by mGluR1 is required for mGluR-LTD

**DOI:** 10.1101/2021.09.14.460352

**Authors:** Luisa Speranza, Yanis Inglebert, Claudia De Sanctis, Pei You Wu, Magdalena Kalinowska, R. Anne McKinney, Anna Francesconi

**Affiliations:** Dominick P. Purpura Department of Neuroscience, Albert Einstein College of Medicine, New York, NY 10461 US; Department of Pharmacology & Therapeutics, McGill University, Montreal, QC H3G 0B1 CA

**Keywords:** Dendritic spines, Synaptopodin, spine apparatus, metabotropic glutamate receptors, mGluR1, mGluR-LTD, lysosome, protein turnover

## Abstract

Dendritic spines, actin-rich protrusions forming the postsynaptic sites of excitatory synapses, undergo activity-dependent molecular and structural remodeling. Activation of group 1 metabotropic glutamate receptors - mGluR1 and mGluR5 - by synaptic or pharmacological stimulation, induces LTD but whether this is accompanied with spine elimination remains unresolved. A subset of telencephalic mushroom spines contains the spine apparatus (SA), an enigmatic organelle composed of stacks of smooth endoplasmic reticulum, whose formation depends on the expression of the actin-bundling protein Synaptopodin. Allocation of Synaptopodin to spines appears governed by cell-intrinsic mechanisms as the relative frequency of spines harboring Synaptopodin is conserved *in vivo* and *in vitro*. Here we show that expression of Synaptopodin/SA in spines is required for induction of mGluR-LTD at Schaffer collateral-CA1 synapses. Post-mGluR-LTD, mushroom spines lacking Synaptopodin/SA are selectively lost whereas spines harboring it are preserved, a process dependent on activation of mGluR1 but not mGluR5. Mechanistically, we find that mGluR1 supports physical retention of Synaptopodin within excitatory spine synapses during LTD while triggering lysosome-dependent degradation of the protein residing in dendritic shafts. Together, these results reveal a cellular mechanism, dependent on mGluR1, which enables selective preservation of stronger spines containing Synaptopodin/SA while eliminating weaker ones and potentially countering spurious strengthening by *de novo* recruitment of Synaptopodin. Overall our results identify spines with Synaptopodin/SA as the locus of mGluR-LTD and underscore the importance of the molecular microanatomy of spines in synaptic plasticity.

## Introduction

Dendritic spines, small protrusions on excitatory neurons, serve as the sites for structural changes during long-term changes in synaptic strength. The mechanisms of long-term potentiation (LTP) associated with spine outgrowth and enlargement (Harris, 2020) are well understood, whereas the occurrence of structural changes in spines after long-term depression (LTD) is more controversial and the underlying mechanisms not fully understood (Stein & Zito, 2019). In some incidences spines are eliminated or reduced in size after LTD but in other cases synaptic weakening can occur independently of spine shrinkage (Thomazeau et al, 2020; Wang et al, 2007). This complexity is compounded by the heterogeneous microanatomy of spines (Berry & Nedivi, 2017), with some marked by transient or enduring presence of smooth endoplasmic reticulum (sER) (Perez-Alvarez et al, 2020; Spacek & Harris, 1997). In telencephalic regions, a subset of large spines contain a spine apparatus (SA) (Spacek, 1985; Spacek & Harris, 1997), a poorly understood neuron-specific sER organelle that supports local Ca^2+^ storage (Fifková et al, 1983; Korkotian et al, 2014). The SA, which is present in ∼10-20% of adult hippocampal and cortical mature spines (Spacek & Harris, 1997), is made of stacks of sER tubules intercalated with F-actin and Synaptopodin (Synpo), an F-actin-bundling protein (Asanuma et al, 2005; Deller et al, 2000). Synpo is necessary and sufficient for the formation and maintenance of the SA, as shown by absence of the organelle in mice lacking Synpo (Deller et al, 2003). At present, the contribution of the SA to forms of activity-dependent synaptic plasticity is incompletely understood and the mechanisms underlying enduring presence of Synpo in subsets of dendritic spines remain unclear.

Activation of group 1 metabotropic glutamate receptors (Gp1 mGluRs) – mGluR1 and mGluR5 - by synaptic or pharmacological stimulation, induces LTD at Schaffer collateral-CA1 synapses (mGluR-LTD) (Lüscher & Huber, 2010), a form of plasticity altered in many neurodevelopmental disorders (Huber et al, 2002; Sahin & Sur, 2015). It was further shown that the capacity of Gp1 mGluRs to depress synaptic transmission relies on sER (Holbro et al, 2009), but whether presence of Synpo is needed for mGluR-LTD to occur is untested. Gp1 mGluRs are critical to circuit refinement during development and their activation was shown to be required for activity-dependent shrinkage of large spines (Oh et al, 2013) and heterosynaptic structural plasticity (Oh et al, 2015). Nevertheless, the impact of mGluR-LTD on spine elimination was recently questioned, since contrasting observations have reported either spine loss induced by mGluR-LTD (Ramiro-Cortés & Israely, 2013; Wiegert & Oertner, 2013) or lack of effect on spine density (Thomazeau et al, 2020). Moreover, whether spines containing the SA undergo structural remodeling during mGluR-LTD is unknown.

Here we show that expression of Synpo/SA in spines is required for induction of mGluR-LTD at hippocampal Schaffer collateral-CA1 synapses. We find that mGluR-LTD induces selective loss of mushroom spines that do not contain Synpo, an effect mediated by mGluR1. We further report that mGluR-LTD triggers selective degradation of Synpo in dendritic shafts *via* the lysosomal pathway, whereas Synpo at spine synapses is stabilized through association with mGluR1. Together, these findings uncover a novel cellular mechanism dependent on mGluR1 that enables selective preservation of spines containing the SA while eliminating weaker ones.

## Materials and Methods

### ANIMALS

All procedures were according to protocols approved by the Albert Einstein College of Medicine, in accordance with the Guide for the Care and Use of Laboratory Animals by the United States PHS or the guidelines of the Canadian Council on Animal Care and the McGill University Comparative Medicine and Animal Resources animal handling protocols 5057. Sprague Dawley rats were used for primary neuronal cultures. *Grm1* knockout mice (*Grm1*^KO^ (Mende et al, 2016)) and *Synp*o knockout mice (Deller et al, 2003) crossed with L15 GFP-expressing mice used as wild type (Verbich et al, 2016) as previously described. Mice of both sexes were used for experiments, fed *ad libitum* and housed with a 12h light/dark cycle.

### ELECTROPHYSIOLOGY

Hippocampal slices were obtained from P 30 to 40 old Synpo KO or L15 mice. Mice were deeply anaesthetized with isoflurane and killed by decapitation. Slices (400 µm) were cut on a vibratome (Leica VT1200S) in an sucrose-based solution containing (in mM) 280 sucrose, 26 NaHCO_3_, 10 Glucose, 1.3 KCl, 1 CaCl_2_ and 10 MgCl_2_ and were transferred at 32°C in regular artificial cerebrospinal fluid (ACSF) containing (in mM) 124 NaCl, 5 KCl, 1.25 NaH2PO4, 2 MgSO4, 26 NaHCO3, 2 CaCl2, and 10 glucose saturated with 95% O2/5% CO2 (pH 7.3, 300 mOsm) for 15 minutes before resting at room temperature for 1 h in oxygenated (95% O2/5% CO2) ACSF. For recording, slices were transferred to a temperature-controlled (32°C) chamber with oxygenated ACSF. To assess mGluR-LTD, slices were placed into a heated (31-32°C) recording chamber of an upright microscope (DM LFSA Microsystems, Heidelberg, Germany) and perfused continuously with regular ACSF. Field excitatory postsynaptic potentials (fEPSPs) were recorded in the *stratum radiatum* of the CA1 region by using glass microelectrodes filled with 3M NaCl. GABA_A_ receptor-mediated inhibition was blocked with 100 µM picrotoxin and the area CA1 was surgically isolated from CA3 to avoid epileptiform activity. fEPSPs were elicited at 0.1 Hz by a digital stimulator that fed by a stimulation isolator unit. All data analyses were performed with custom written software in Igor Pro 8 (Wavemetrics). fEPSP slope was measured as an index of synaptic strength.

### NEURONAL CULTURES AND DRUG TREATMENTS

Hippocampi and cortices from newborn rat pups dissected in Ca^2+/^Mg^2+^-free HBSS were digested with 0.25% trypsin and DNAse (2000 U/ml) for 20 min at 37°C and mechanically triturated. Viable cells, determined by trypan blue dye exclusion, were plated at 8×10^4^/1.13cm^2^ on poly-L-lysine-coated cover glasses and at 2.5×10^5^/1.9cm^2^ in multi-well plates. Cells were maintained at 37°C, 5% CO_2_ in serum-free Neurobasal A medium with 2% B-27 supplement, 2 mM GlutaMax (from Invitrogen), penicillin (50 U/ml), streptomycin (50 μm/ml). After 5 days *in vitro*, a mix of 37 mM Uridine and 27 mM 5-Fluoro-2-deoxyuridine was added; half medium was replaced weekly and neurons used at 19-21 days *in vitro*.

For chemical mGluR-LTD, cells were rinsed with pre-warmed medium and treated with 50 µM *S*-DHPG (Tocris Bioscience) or vehicle for 15 min at 37°C: cells were rinsed with pre-warmed medium and incubated with fresh medium for 30 or 120 min at 37°C. Bay 36-7620 or MPEP (Tocris Bioscience) were applied alone for 15 min at 37°C or together with DHPG and added to the fresh medium during recovery. For treatment with MG132 (ApexBio), leupeptin or BafA_1_ (Sigma Aldrich), after rinsing with pre-warmed medium, drugs or respective vehicle were applied alone in fresh medium for 15 min at 37°C or together with DHPG, and added to fresh medium during recovery. For treatment with rapamycin (Cayman Chemical) or cycloheximide (Sigma Aldrich), cells were rinsed with pre-warmed medium and drugs or vehicle applied in fresh medium for the indicated times at 37°C. After treatments, cells were placed on ice and processed for downstream analysis.

### DiIC_18_ LABELING

Cells and tissue sections were labeled with the fluorescent dye 1,1’-Dioctadecyl-3,3,3’,3’-Tetramethylindotricarbocyanine Iodide (DiIC_18_; Molecular Probes, crystals cat. D3911) as described (Cheng et al, 2014). Briefly, cells plated on cover glasses were fixed with 4% paraformaldehyde (PFA) for 10 min and washed with PBS. After PBS aspiration, DiIC_18_ crystals (3-4 crystals/well) were sprinkled over the cells with a 18 gauge needle. To prevent dehydration, PBS was added to the wells and the plate incubated for 10 min on an orbital shaker at RT protected from light. After washing with PBS to remove the crystals, cells were incubated in PBS overnight at RT and washed three times with PBS (5 min per wash) before immunolabeling or mounting on glass slides. For tissue sections, mice were anesthetized with isoflurane and perfused with 4% PFA: the brain was removed and post-fixed with 4% PFA overnight at 4°C. After three 5 min washes with PBS, 150 or 300 *μ*m-thick coronal sections were cut with a vibratome (Leica VT1000S). The tissue was gently unfolded with a paintbrush and crystals of DiIC_18_ applied with a needle onto the regions of interest (parietal and prefrontal cortex). Labeled tissue sections were incubated at 4°C in PBS, protected from light, for 7 to 10 days before image acquisition.

### IMMUNOFLUORESCENCE

Cells were rinsed in PBS for 2 min and fixed in 4% PFA for 10 min at RT. After permeabilization with 0.1% Triton X-100 in PBS for 15 min, cells were incubated for 1 h at RT with blocking solution of either 5% bovine serum albumin (BSA) or 5% normal serum. Primary antibodies in blocking solution were applied overnight at 4°C. After three washes with PBS for 3 minutes, cells were incubated for 1-2 h at RT with fluorophore-conjugated secondary antibodies, washed three times with PBS and mounted on glass slides with Prolong (Invitrogen). To label surface AMPARs, cells were rinsed two times with pre-warmed medium and incubated for 30 min at 37°C with rabbit anti-GluA1 (1/150; Calbiochem/Millipore, RRID: AB_564636) in culture medium. After incubation with anti-GluA1, cells were treated with 50 µM DHPG or vehicle for 15 min at 37°C, rinsed with pre-warmed medium, and incubated with fresh medium for 30 min at 37°C. Cells were then washed with PBS containing 1 mM MgCl_2_, 0.1 mM CaCl_2_ and fixed with 4% PFA for 10 min. After fixation, cells were washed with PBS (3x 5 min), incubated with 5% BSA for 1 h at RT and with secondary fluorescent antibody for 1 h at RT. After washing with PBS, cells were permeabilized, blocked and incubated with primary antibodies as above. The following antibodies were used: rabbit anti-Synaptopodin (1/500; Synaptic Systems, RRID: AB_887825), guinea pig anti-Synaptopodin (1/500; Synaptic Systems, RRID: AB_10549419) chicken anti-MAP2 (1/500; PhosphoSolutions, RRID: AB_2138173), rabbit anti-mGluR1 (1/2,000; RRID: AB_2571735). The following secondary antibodies were used: donkey anti-mouse or rabbit conjugated to Alexa Fluor 488 and Alexa Fluor 647, goat anti-guinea pig Alexa Fluor 488 (Invitrogen), donkey anti-chicken IgY conjugated to Alexa Fluor 647 and Aminomethylcoumarin Acetate (Jackson ImmunoResearch Labs)

### MICROSCOPY

Wide field fluorescence imaging was carried out with an Olympus IX81 inverted microscope equipped with digital CCD ORCA-R2 camera (Hamamatsu) using 40x (N.A.=1.3) or 60x (N.A.=1.35) oil objectives. Confocal images were acquired with a Zeiss LSM880 Airyscan using a Plan-Apochromat 63x (N.A.=1.4) oil immersion objective; a 561 nm diode pumped solid-state laser (DPSS 561-10) was used to visualize DiIC_18_. Images at 1024×1024 pixel resolution were acquired with scan speed set at 6 and pinhole configured to 1 Airy unit for each channel. Stacks of images were acquired with a 0.5 *μ*m *Z* step and reconstructed with the Fiji’s Stacks *Z* Project function to generate *Z*-stack projections of the maximum intensity.

### IMAGE ANALYSIS

Analysis was carried out blind to treatment/genotype using the image-processing platform Fiji (Schindelin et al, 2012). To quantify Synpo clusters, a mask of the outline of neurons was generated using MAP2 signal overlay. After background subtraction, individual clusters within the masked region were measured by automated count using the Analyze Particles function on thresholded images. For spine analysis, dendrites and dendritic protrusions were outlined and measured with the segmented line tool. Proximal dendritic segments (∼100 *μ*m) were analyzed in both primary and secondary dendrites. Spine density is expressed as number of spines per 10 *μ*m of dendritic length. To classify dendritic spines, the length (*l*) head width (*h*) and neck width (*n*) were manually traced for individual dendritic protrusions. Spine length was measured from the emerging point on the shaft to the tip of spine head, and head diameter was measured at the point of maximum width. Dendritic protrusions were classified according to established morphological criteria (Harris et al, 1992; Zagrebelsky et al, 2005) and defined as mushroom spines when *h* » *n* (*h/n* > 1.5) and thin spines when *l* > 1 *μ*m and *h/n* ≤ 1.5. Stubby spines and filopodia were rarely observed in the mature neurons used in the study and not included in the analysis.

### WESTERN BLOT AND IMMUNOPRECIPITATION

Drug-treated cortical neurons were placed on ice, rinsed with ice-cold PBS and lysed in buffer of 20 mM MOPS (pH 7.2), 2 mM EGTA, 5 mM EDTA, 1% Triton X-100 supplemented with a cocktail of protease and phosphatase inhibitors. Cells were incubated on ice for 5 min, scraped off the wells, and collected by centrifugation at 21,000 x *g* at 4°C for 15 min. Equal amount of proteins were resolved by SDS-PAGE and processed by Western blot assay per standard protocols using horseradish peroxidase-conjugated secondary antibodies and ECL for detection on film or with an Azure c600 imaging system (Azure Biosystems). The following primary antibodies were used: rabbit anti-Synaptopodin (1/1000), goat anti-Synaptopodin (1/200; Santa Cruz Biotech, RRID: AB_2201166), rabbit mGluR1 (1/500, Alomone labs, RRID:AB_2039984; 1/10,000 RRID:AB_2571736), mouse anti-PSD95 (1/400; Antibodies Inc., RRID: AB_2292909), mouse anti-*γ*-Tubulin (1/2,500; Sigma Aldrich, RRID: AB_477584).

For immunoprecipitation, brain cortices of adult mice were homogenized on ice in a buffer of 20 mM Tris-HCl, 5 mM EDTA, 100 mM NaCl, 1% Triton X-100, 0.5% sodium deoxycholate (pH 7.4; 10 *μ*l/mg tissue), supplemented with cocktails of protease and phosphatase inhibitors. The tissue was manually homogenized on ice, centrifuged at 13,000 x *g* for 20 min at 4°C and supernatant collected. Equal amounts of protein were incubated overnight at 4°C with primary antibodies (rabbit anti-Synaptopodin 3 µl/mg; goat anti-Synaptopodin 2 µg) or control IgG coupled to Protein G-coupled magnetic beads (Dynabeads, Life Technologies). Unbound material was removed and the beads washed three times with homogenization buffer, three times with PBS/0.1% Triton X-100 and one time with PBS before elution in denaturing sample buffer at 95°C for 5 min.

### STATISTICAL ANALYSIS

Unless noted, values were imported into Prism 8.1 (GraphPad) for the generation of graphs and statistical tests. Data are reported as mean ± SEM unless indicated: normality and data distribution were assessed with the Kolmogorov-Smirnov test. Two-tailed Student *t*-test was used to compare two groups and Mann-Whitney test used for non-parametric analysis of ranks when appropriate. One-way ANOVA with Tukey’s post-hoc test was used to compare multiple groups. Gardner-Altman estimation plots used to display spine data and the two-sided permutation *t-*test, were generated with the web application estimationstats.com (Ho et al, 2019). *P* value < 0.05 was considered significant in all statistical comparisons.

## Results

### Spines with stable Synpo are spared from elimination and required for mGluR-LTD

Dendritic spines are the sites of excitatory synapses and their structural and functional properties are modified in response to activity patterns that produce long-term changes in synaptic transmission. Low frequency stimulation was shown to selectively induce depression of evoked EPSCs in dendritic spines containing sER, an effect that was blocked by Gp1 mGluR antagonists (Holbro et al, 2009). Although this finding suggested that Gp1 mGluR-induced synaptic depression could be compartmentalized to spines endowed with ER, whether the presence of Synpo is required for mGluR-LTD and if it plays a role in spine elimination after mGluR-LTD is currently unknown. To address this question, we used DHPG (*RS*-DHPG; 100 µM, 5 min) to induce mGluR-LTD at stimulated Schaffer collateral (SC)-CA1 synapses (**Fig. 1*A***) in hippocampal slices from WT and *Synpo* knockout mice (*Synpo*^KO^) which lack the SA (Deller et al, 2003). In WT, DHPG induced rapid and enduring depression of evoked field potential (fEPSP) slope values compared to baseline, as measured 35 min post-DHPG (fEPSPs to 69% ± 6 of baseline, *n* = 9; **Fig. 1*B,C***). In contrast, in *Synpo*^KO^ mice the capacity of DHPG to produce LTD was drastically impaired (depression of fEPSPs to 87 ± 3% of baseline, *n* = 9; **Fig. 1*B,C***). Thus this finding indicates that expression of Synpo and presence of the SA is required for efficient mGluR-LTD at SC-CA1 synapses.

**Figure 1.**
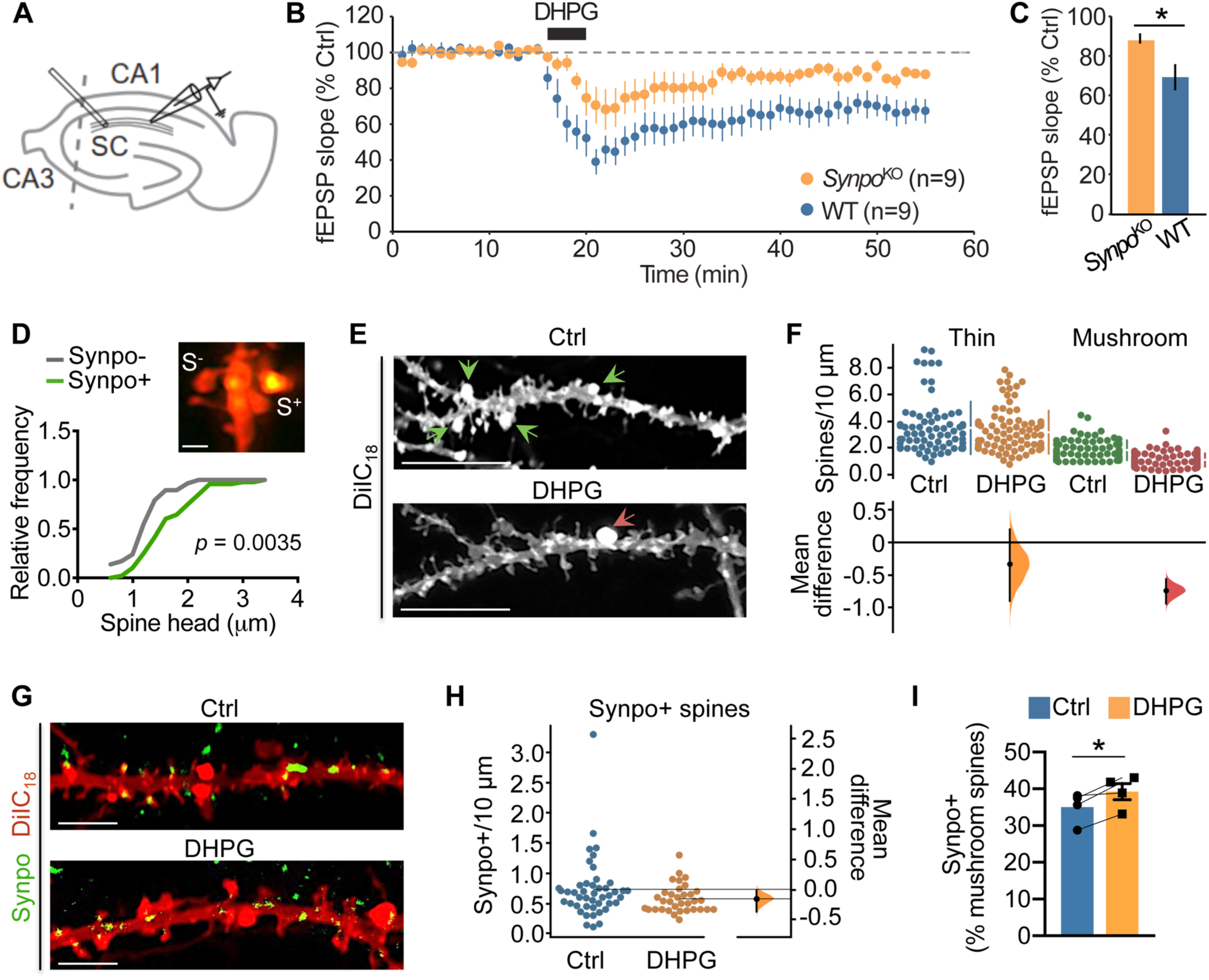
Spines with stable Synpo are spared from elimination and required for mGluR-LTD. A. Illustration of the approximate locations of stimulating and recording electrodes in hippocampal slices to measure mGluR-LTD at Schaffer Collateral (SC) to CA1 synapses. B. Time-course of synaptic changes induced by bath application of 100 µM DHPG during 5 minutes in WT (n=9 slices, N=4 mice) and *Synpo*^KO^ mice (n=9 slices, N=5 mice). C. Quantification of average mGluR-LTD in the last 10 min of the recording: mean ± SEM *Synpo*^KO^ 87% ± 3, WT 69% ± 6, n=9 slices, *p*<0.05, Mann-Whitney test. D. Characterization of mushroom spines with or without Synpo. Representative image (Airyscan) of spines in hippocampal neurons labeled with DiIC_18_ and anti-Synpo. S+, spines with Synpo, S-, spines without Synpo: scale bar 0.5 *μ*m. Relative frequency distribution of spine head dimensions. Synpo- (*n*=29), Synpo+ (*n*=48), Kolmogorov-Smirnov test. E. Images of DiIC_18_-labeled neurons treated with vehicle (Ctrl) or mGluR-LTD (DHPG; 30 min recovery); arrowheads indicate mushroom spines. Scale bars 10 *μ*m. F. Quantification of thin and mushroom spine density per 10 *μ*m. Mean ± SD, thin spines Ctrl 3.5 ± 2.0 (n=74 dendritic branches), DHPG 3.2 ± 1.6 (n=91), N=36 cells per group, *p*=0.246 two-sided permutation *t*-test; mushroom spines, Ctrl 1.9 ± 0.74 (n=78), DHPG 1.1 ± 0.50 (n=109), N=45 cells, *p*<0.0001. Gardner-Altman estimation plot: mean difference (dot, bottom panel) is plotted on a floating axis as a bootstrap sampling distribution, vertical error bars indicate the 95% confidence interval. G. Representative images of hippocampal neurons treated with vehicle (Ctrl) or DHPG (30 min recovery) and labeled with DiIC_18_ and anti-Synpo antibody; scale bars, 4 *μ*m. H. Quantification of Synpo+ spines in neurons treated with vehicle (Ctrl) or DHPG from images like those in (g); spines/10 *μ*m, mean ± SD, Ctrl 0.73 ± 0.51 (n=45), DHPG 0.58 ± 0.23 (n=34), N=30 cells per group, *p*=0.096 two-sided permutation t-test. I. Percentage of Synpo+ spines relative to total mushroom spines in neurons treated with vehicle (Ctrl) or mGluR-LTD (30 min): mean ± SE from N=4 experiments, *p*=0.04, paired t-test.

NMDAR-dependent long-term depression was shown to causes spine shrinkage and elimination (Stein & Zito, 2019), but whether mGluR-LTD is accompanied with spine elimination remains controversial. Both spine shrinkage and elimination (Ramiro-Cortés & Israely, 2013) and, conversely, absence of changes in structural plasticity and spine density (Thomazeau et al, 2020) were reported in response to Gp1 mGluR activation. In addition, the specific impact of mGluR-LTD on spines containing Synpo/SA is unknown. Synpo is expressed in spines, starting around postnatal day 12, and is most abundant in the adult brain (Czarnecki et al, 2005). Remarkably, its developmental regulation is mirrored in primary neurons (Konietzny et al, 2019), where Synpo is detected at ∼1 week of differentiation *in vitro* and increases over time, peaking at ∼3 weeks. Congruent with its intrinsic association with the SA, Synpo occupies the neck and head of mushroom spines (**Fig. 1*D***), ∼50% of which are of size comparable to mushroom spines lacking Synpo so that they are not distinguishable based on morphology alone (**Fig. 1*D***). To examine the impact of mGluR-LTD on spine structural plasticity and elimination, we used sparse labeling with the fluorescent lipophilic dye DiIC_18_ and high-resolution microscopy to survey density and structural properties of spines under physiological conditions that preserve actin dynamics (McKinney, 2010; Roelandse & Matus, 2004) and rate of protein synthesis (Cooke et al, 2019), both crucial for structural and functional plasticity (Cingolani & Goda, 2008; Huber et al, 2000). Widespread chemical mGluR-LTD was produced by transient stimulation with *S*-DHPG (50 µM, 15 min) followed by recovery for 30 or 120 min, corresponding to early and late times post-induction when LTD is fully expressed. In concurrence with previous data (Snyder et al, 2001), chemical mGluR-LTD (mGluR-LTD hereafter) decreased the surface abundance of the GluA1 subunit of AMPARs (surface GluA1 puncta per cell, fold steady state: mean ± SE, steady state 1.0 ± 10.039 n=13, vehicle 0.97 ± 0.052 n=13, DHPG 0.81 ± 0.051 n=11, vehicle *vs*. DHPG *p* = 0.019, ANOVA with Tukey’s post-test) attesting to the effectiveness of the LTD protocol. We found that mGluR-LTD resulted in a net decrease in the overall density of dendritic spines at 30 min post-treatment (total protrusions/10µm, mean±SD vehicle 3.7±1.1 n=54 dendritic branches *vs*. mGluR-LTD 2.7±0.76 n=78 from N=30 neurons per group; *p*<0.001, Mann-Whitney test). Categorization of spines based on established morphological criteria indicated that mGluR-LTD produced net loss of the synaptically stronger mushroom spines whereas the density of thin spines was unaffected (**Fig. 1*E,F***).

The finding that mGluR-LTD is impaired in *Synpo*^-/-^ mice suggests that spines containing Synpo are a primary *locus* of the plasticity. Therefore we next tested whether the observed net loss of mushroom spines induced by mGluR-LTD might arise from elimination of spines containing Synpo. Visualization of endogenous Synpo by immunolabeling of neurons filled with DiIC_18_ shows that, as in the hippocampus where ∼10% of total spines contain Synpo/SA, in primary hippocampal neurons ∼13% of all dendritic protrusions contain Synpo (mean±SD, 13±4.6% n=17) accounting for ∼30% of mushroom spines (mean±SD, 32±13% n=44). Surprisingly, we found that despite net loss of mushroom spines, the density of Synpo-containing spines did not significantly differ after mGluR-LTD compared to control (**Fig. 1*G,H***) so that the relative frequency of Synpo-positive *vs*. total mushroom spines was modestly increased by mGluR-LTD (**Fig. 1*I***). Both mGluR1 and mGluR5 contribute to induction of mGluR-LTD by DHPG but only selective inhibition of mGluR1 was shown to revert LTD and AMPAR internalization (Volk et al, 2006) suggesting distinct receptor functions. To test whether both mGluR1 and mGluR5 play a role in net loss of Synpo-lacking mushroom spines, mGluR-LTD was induced in the presence of either Bay 36-7620 (Bay; 10 µM) or 2-Methyl-6-(phenylethynyl) pyridine (MPEP; 10 µM), selective inverse agonists for mGluR1 and mGluR5, respectively. We found that whereas Bay halted mGluR-LTD-induced net loss of mushroom spines (**Fig. 2*A,B***), MPEP had no significant effect (**Fig. 2*C,D***). Collectively, these findings indicate that mGluR-LTD induces selective loss of mushroom spines that do not contain Synpo whereas those in which Synpo/SA is present are spared, a process that is dependent on mGluR1 activation during mGluR-LTD.

**Figure 2.**
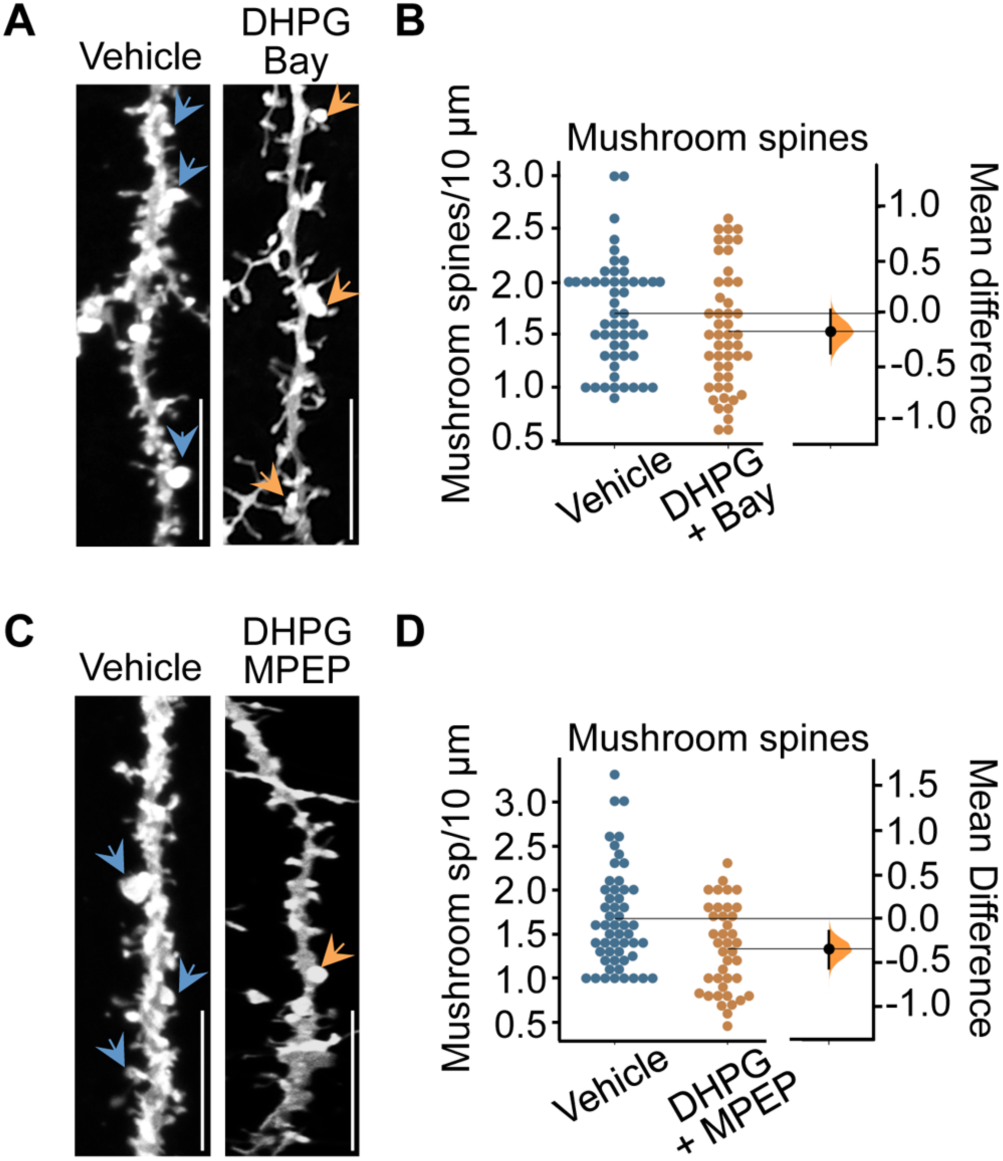
mGluR-LTD induces loss of mushroom spines through activation of mGluR1. A. Representative images of hippocampal neurons treated with vehicle or DHPG in the presence of Bay (30 min recovery) and labeled with DiIC_18_; arrowheads indicate mushroom spines, scale bars, 10 µm. B. Quantification of mushroom spine density from images like those in (a): mean ± SD, vehicle 1.7 ± 0.50 (n=55), DHPG+Bay 1.5 ± 0.57 (n=49), N=25 cells per group, *p*=0.1044, two-sided permutation *t* test. C. Representative images of hippocampal neurons treated with vehicle or DHPG in the presence of MPEP (30 min recovery) and labeled with DiIC_18_; scale bars, 10 µm. D. Quantification of mushroom spine density from images like those in (c): mean ± SD, vehicle 1.7 ± 0.57 (n=53), DHPG+MPEP 1.3 ± 0.49 (n=40) from 3 experiments, *p*=0.0032, two-sided permutation *t* test.

### mGluR-LTD induces compartment-specific degradation of Synpo

Thus far our findings indicate that the proportion of mature mushroom spines tagged by Synpo remains constant in response to mGluR-LTD, suggesting that overall only mushroom spines in which Synpo is absent are lost. Hence whereas spines tagged by Synpo and SA remain stable, those ‘untagged’ are eliminated. If so, how can Synpo ‘tagging’ be restricted to subsets of spines? One possibility is that Synpo expression and availability for recruitment to spines could be constrained. In addition to being localized in spines, Synpo is present in dendritic shafts where it appears in discrete clusters and occasional tubule-like structures (Konietzny et al, 2019)(Deller et al, 2000). Various synaptic activity patterns were previously shown to regulate Synpo expression, including LTP (Yamazaki et al, 2001) and denervation-induced synaptic strengthening (Vlachos et al, 2013) that increase Synpo biosynthesis or enlargement of Synpo clusters, respectively. Homeostatic synaptic downscaling (Dörrbaum et al, 2020) in elevated activity induces instead a decrease in Synpo abundance.

Next we examined the impact of mGluR-LTD on Synpo expression using immunofluorescence in hippocampal neurons to visualize dendritic Synpo clusters and immunoblot in cortical neurons to monitor changes in Synpo abundance. In dendrites visualized by MAP2, mGluR-LTD reduced the density of Synpo clusters per cell within 30 min of stimulation, an effect that persisted at 120 min post-DHPG (**Fig. 3*A,B***). In line with results by immunofluorescence, the total abundance of Synpo protein was similarly reduced in cortical neurons after mGluR-LTD (**Fig. 3*C,D***). We next tested if both mGluR1 and mGluR5 contributed to the loss of dendritic Synpo by inducing mGluR-LTD in the presence of either Bay or MPEP, respectively. In the presence of Bay, mGluR-LTD failed to decrease Synpo clusters whereas Bay alone had no significant effect (**Fig. 3*E,F***). In contrast, in the presence of MPEP, mGluR-LTD effectively reduced dendritic Synpo clusters and application of MPEP alone caused a modest decrease in Synpo (**Fig. 3*G,H***). We reasoned that the effect of MPEP may be linked to an ability of mGluR5 to promote Synpo synthesis, potentially *via* signaling to mTOR (Richter & Klann, 2009). We therefore examined Synpo expression and turnover in neurons treated with the mTOR inhibitor rapamycin (**Fig. 4*A,B*)** or the protein synthesis inhibitor cycloheximide, respectively (**Fig. 4*C-F***). We found that Synpo abundance decreased under both conditions, suggesting that constitutive basal biosynthesis of Synpo is mTOR-dependent (Dörrbaum et al, 2020). Together, these findings indicate that Synpo undergoes constitutive turnover and that mGluR1 activation during mGluR-LTD produces a rapid, selective decrease of Synpo protein present in dendritic shafts.

**Figure 3.**
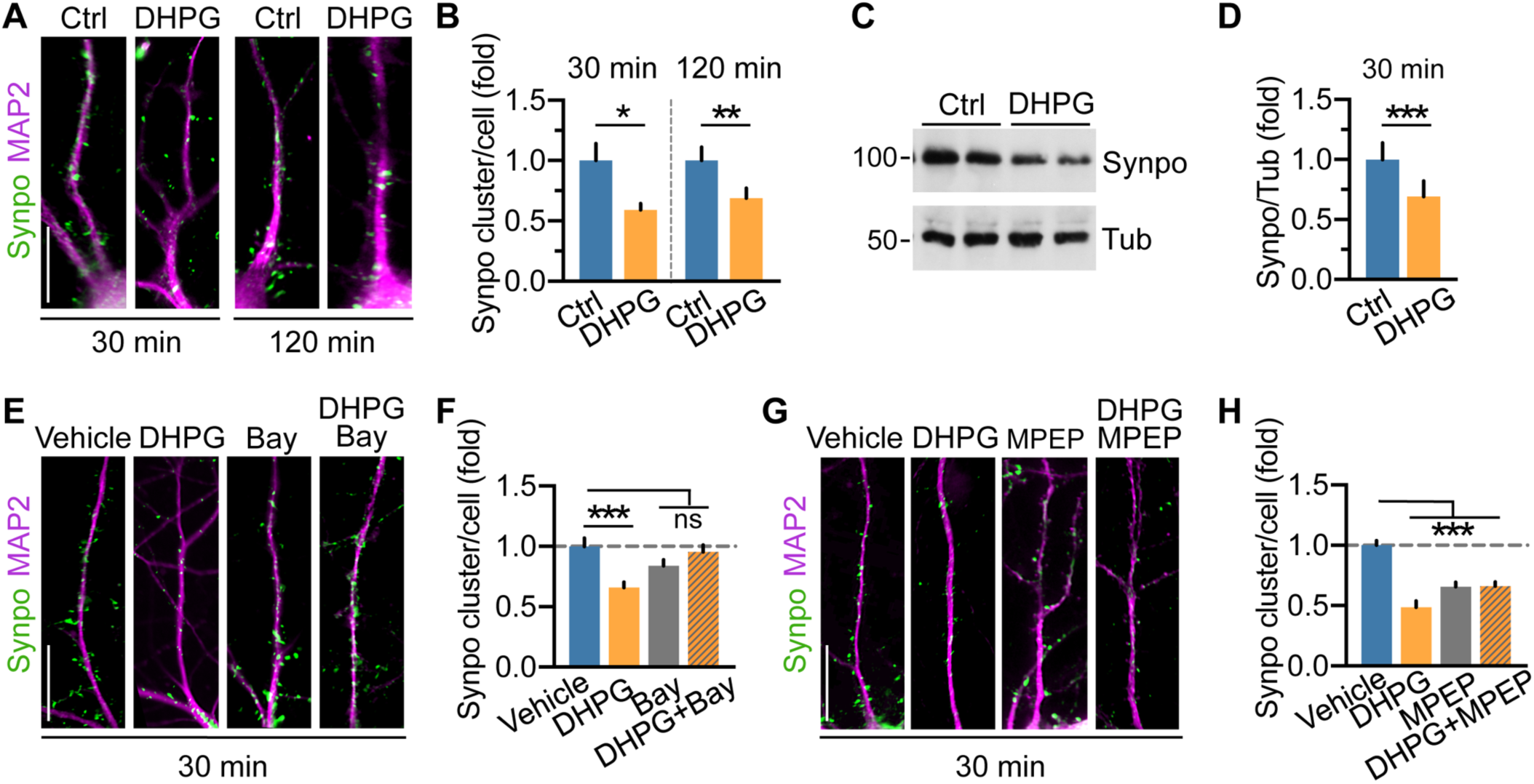
mGluR-LTD induces degradation of Synpo in dendritic shafts. A. Representative images and quantification of immunolabeled Synpo in MAP2-positive dendrites of hippocampal neurons treated with vehicle (Ctrl) or DHPG followed by recovery for 30 or 120 min. Scale bars, 15 µm. B. Quantification of dendritic Synpo clusters per cell (fold vehicle) from images like in (a): mean ± SE 30 min, Ctrl 1.0 ± 0.14, DHPG 0.59 ± 0.053, n=71 cells per group from N=6 experiments, (*) *p*=0.0106; 120 min, Ctrl 1.0 ± 0.111 (n=44), DHPG 0.69 ± 0.083 (n=45), N=4, (**) *p* = 0.004, paired t-test. C. Representative immunoblot of cortical neurons treated with vehicle (Ctrl) or DHPG followed by 30 min recovery: Tubulin (Tub), loading control. D. Quantification of Synpo (fold Ctrl) from blots as in (c): mean ± SE, Ctrl 1.0 ± 0.14, DHPG 0.69 ± 0.13, N=5, (***) *p* = 0.0002, paired t-test. E. Representative images of Synpo in hippocampal neurons treated with vehicle, Bay or DHPG (in presence/absence of Bay) followed by 30 min recovery. Scale bar 15 µm. F. Quantification of Synpo clusters from images as in (e). Mean ± SE, vehicle 1.0 ± 0.069 (n=17), DHPG 0.66 ± 0.044 (n=28), Bay 0.84 ± 0.049 (n=26), DHPG+Bay 0.95 ± 0.058 (n=18) from 2 experiments, (***) *p*<0.001, (ns) *p*>0.1, ANOVA with Tukey’s post-test. G. Representative images of Synpo in hippocampal neurons treated with vehicle, MPEP or DHPG in presence/absence of MPEP followed by 30 min recovery. Scale bar 15 µm. H. Quantification of Synpo clusters from images as in (g). Mean ± SE, vehicle 1.0 ± 0.039 (n=29), DHPG 0.49 ± 0.052 (n=16), MPEP 0.66 ± 0.038 (n=32), DHPG+MPEP 0.66 ± 0.035 (n=31); from 4 experiments, (***) *p*<0.001, ANOVA with Tukey’s post-test.

**Figure 4.**
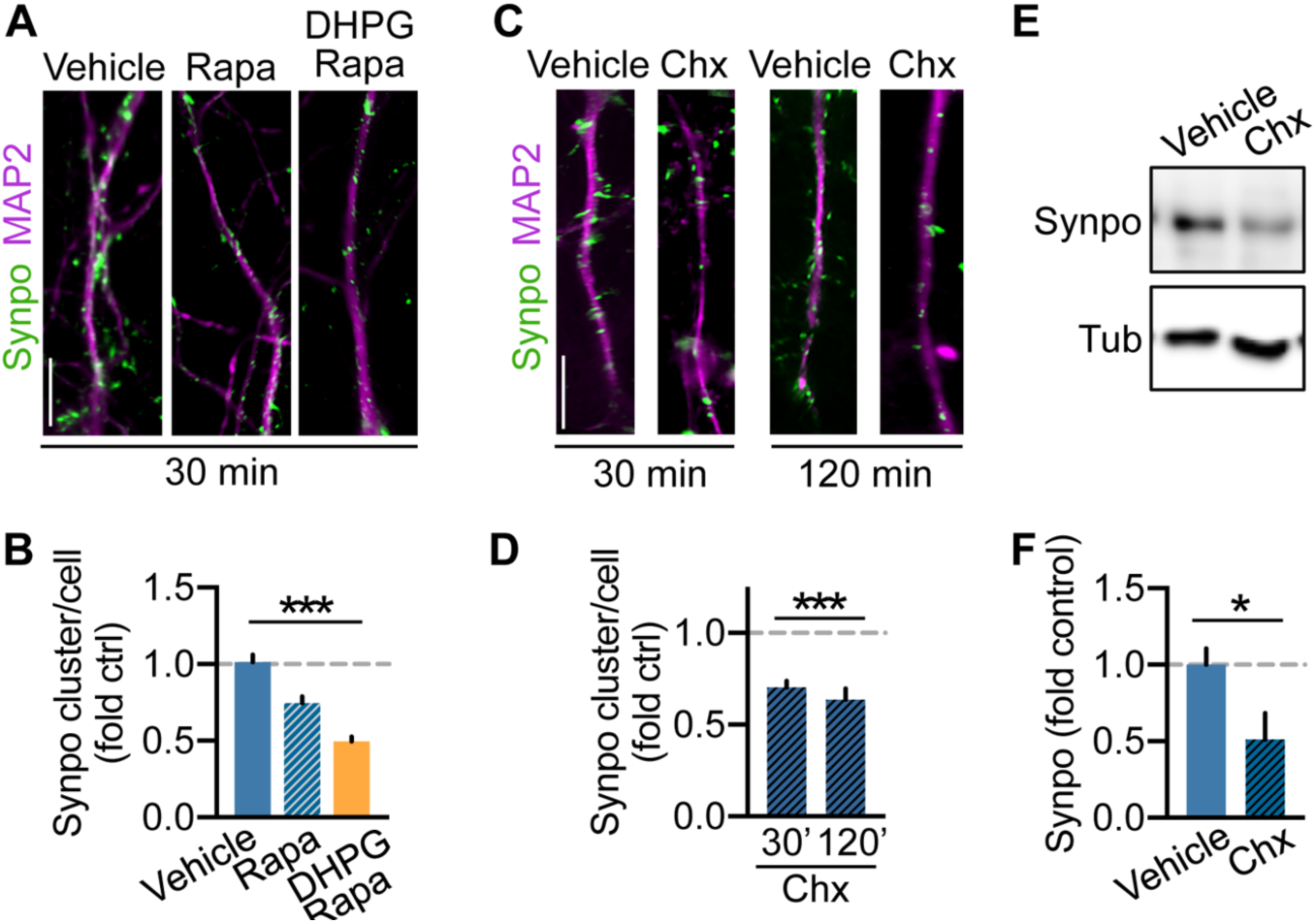
Basal expression of Synpo is regulated by mTOR and ongoing protein synthesis. A. Representative images of immunolabeled Synpo in MAP2-positive dendrites of hippocampal neurons treated with vehicle, rapamycin (20 nM) or DHPG in the presence of rapamycin (30 min recovery); scale bars, 10 µm. B. Quantification of Synpo clusters per cell (fold control vehicle) from images as in (a). Mean ± SE vehicle 1.0 ± 0.048 (n=23), rapamycin 0.75 ± 0.042 (n=40), DHPG+rapamycin 0.50 ± 0.031 (n=30), 2 experiments; (***) *p*<0.001, ANOVA with Tukey’s post-test. C. Representative images of dendritic Synpo in hippocampal neurons treated with vehicle or Chx (50 µM) for 30 min and 120 min; scale bar 10 µm. D.Quantification of Synpo clusters (fold vehicle) from images as in (b). Mean ± SE 30 min, vehicle 1.0 ± 0.032 (n=13), Chx 0.70 ± 0.036 (n=18); 120 min vehicle 1.0 ± 0.033 (n=10), Chx 0.64 ± 0.060 (n=14), from 2 experiments, (***) *p*<0.001, ANOVA with Tukey’s post-test. E.Representative immunoblot of cortical neurons treated with vehicle or Chx for 120 min. F.Quantification of total Synpo from blots as in (e): mean ± SE, vehicle 1.0 ± 0.11 (n=5), Chx 0.51 ± 0.17 (n=5), 2 experiments, (*) *p*=0.042, two-tailed unpaired t-test.

### mGluR-LTD promotes Synpo degradation *via* lysosomal proteolysis

Synpo is a natively unfolded protein (Chalovich & Schroeter, 2010), the primary structure of which includes proline-rich regions that mark unstable proteins. Natively unfolded proteins tend to form aggregates that can be cleared *via* autophagy through lysosomal digestion (Lamark & Johansen, 2012). To test if lysosomal proteolysis contributes to Synpo turnover, hippocampal neurons were treated with leupeptin (Leu; 200 µM) - an inhibitor of lysosomal cysteine, serine and threonine peptidases - for either 30 min or 120 min. In these conditions, the density of Synpo clusters was not significantly altered after 30 min but increased over 120 min (**Fig. 5*A,B***) concordant with a leupeptin-dependent increase in total Synpo in cortical neurons (**Fig. 5*C,D***). Induction of mGluR-LTD in the presence of leupeptin failed to reduce the number of dendritic Synpo clusters (30 min and 120 min; **Fig. 5*A,B***) and to decrease total Synpo protein abundance (**Fig. 5*C,D***). To confirm a role of lysosomal proteolysis in this process, we used Bafilomycin A_1_ (BafA_1_; 100 nM), a V-ATPase inhibitor that prevents lysosome acidification and blocks autophagosome–lysosome fusion (Mauvezin et al, 2015). In the presence of BafA_1_, mGluR-LTD failed to decrease dendritic Synpo (**Fig. 5*E,F***) but did produce a modest enlargement of individual Synpo clusters compared to vehicle or mGluR-LTD alone (**Fig. 5*G***), suggesting accumulation of Synpo in disabled lysosomes.

**Figure 5.**
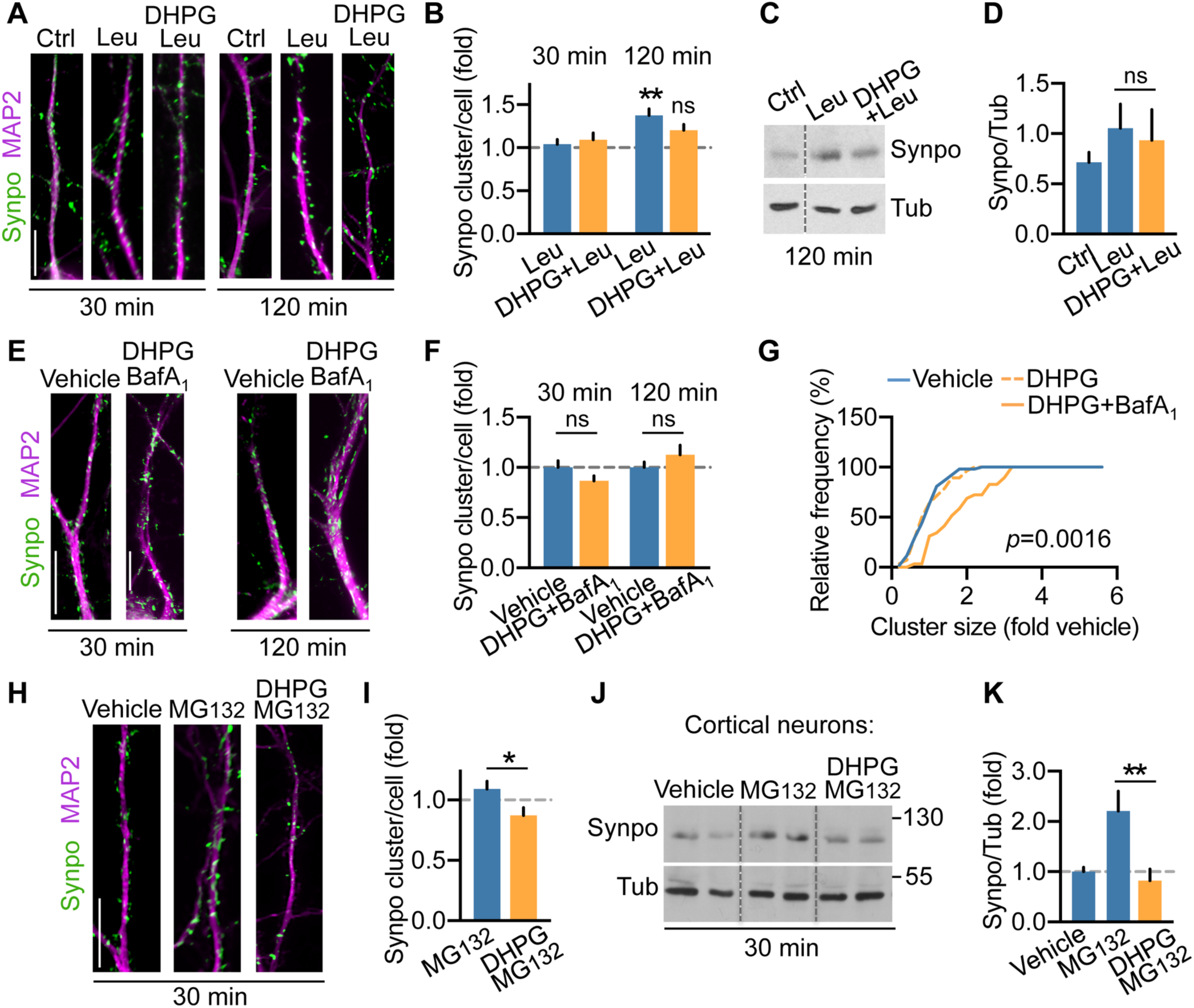
mGluR-LTD promotes Synpo degradation via lysosomal proteolysis. A. Representative images of Synpo in hippocampal neurons treated with vehicle (Ctrl), leupeptin (Leu), DHPG in the presence/absence of leupeptin for 30 or 120 min. B. Quantification of Synpo clusters from images as in (a). Mean±SE 30 min, Ctrl 1.0 ± 0.06 (n=30), Leu 1.04 ± 0.054 (n=35), DHPG+Leu 1.1 ± 0.079; 120 min, Ctrl 1.0 ± 0.076 (n=19), Leu 1.4 ± 0.076 (n=36), DHPG+Leu 1.2 ± 0.07 (n=26), 2 experiments; (ns) p=0.22, (**) *p*=0.004, ANOVA, Tukey’s post-test. C. Representative immunoblot of cortical neurons treated with vehicle (Ctrl), Leu, DHPG in the presence of Leu (120 min recovery); Tubulin (Tub), loading control. D. Quantification of blots as in (c). Total Synpo (fold Tub): mean ± SE, Ctrl 0.71 ± 0.05, Leu 1.1 ± 0.12, DHPG+Leu 0.93 ± 0.15, 4 independent determinations, (ns) *p*=0.640, two-tailed unpaired t-test. E. Representative images of Synpo in hippocampal neurons treated with vehicle or DHPG in the presence of BafA1 for 30 or 120 min; scale bar 15 µm. F. Quantification of Synpo clusters from images as in (e). Mean ± SE 30 min, vehicle 1.0 ± 0.066, (n=24), DHPG+BafA_1_ 0.87 ± 0.049 (n=36), 2 experiments, (ns) *p*=0.41; 120 min, vehicle 1.0 ± 0.054, (n=26), DHPG+BafA_1_ 1.13 ± 0.095 (n=20), (ns) *p*=0.56, ANOVA, Tukey’s post-test. G. Distribution of Synpo cluster size (fold vehicle) in cells treated with vehicle (n=53) or DHPG (30 min recovery) in absence (n=28) or presence of BafA_1_ (n=29); Kolmogorov-Smirnov test. H. Representative images of Synpo in neurons treated with vehicle, MG132, DHPG+MG132 followed by recovery in MG132 (30 min): scale bar 15 µm. I. Quantification of Synpo clusters from images as in (h). Mean ± SE, vehicle 1.0 ± 0.083 (n=20), MG132 1.1 ± 0.064 (n=27), DHPG+MG132 0.87 ± 0.064 (n=25), 2 experiments, (*) *p*=0.0196, two-tailed unpaired t-test. J. Representative blot of cortical neurons treated with vehicle, MG132, DHPG+MG132 with recovery in the presence of MG132. K. Quantification of Synpo from blots as in (j): mean ± SE, vehicle 1.0 ± 0.088 (n=10), MG132 2.2 ± 0.39 (n=5), DHPG+MG132 0.82 ± 0.23 (n=7), 3 experiments, (**) *p*=0.001, ANOVA, Tukey’s post-test.

Next, we tested if the proteasome could also participate to DHPG-induced Synpo degradation since proteasome activity was shown to contribute to protein turnover during mGluR-LTD (Citri et al, 2009; Hou et al, 2006; Klein et al, 2015) and homeostatic down-scaling (Dörrbaum et al, 2020). Treatment of hippocampal neurons with the proteasome inhibitor MG132 (10 *μ*M, 30 min) produced a modest (not statistically significant) increase in the density of dendritic Synpo but did not block mGluR-LTD-induced loss of Synpo clusters (**Fig. 5*H,I***). Similarly, incubation with MG132 increased total Synpo abundance in cortical neurons but did not prevent mGluR-LTD-induced degradation **(Fig. 5*J,K***) suggesting that although Synpo turnover is partly mediated by the proteasome, its degradation by mGluR-LTD is proteasome-independent. Altogether, these results indicate that mGluR-LTD induces rapid degradation of dendritic Synpo *via* the lysosomal pathway.

### mGluR1 contributes to Synpo stabilization in dendritic spines

Our findings indicate that mGluR-LTD, primarily through activation of mGluR1, triggers selective degradation of Synpo present in dendritic shafts but not of Synpo targeted to spines that are spared by mGluR-LTD-dependent elimination. These observations raise the question of how Synpo could be stabilized and shielded from degradation in spines. Synpo, an actin-bundling protein, is recruited to spines through association with α–actinins (Asanuma et al, 2005; Kremerskothen et al, 2005), including α–actinin-4, which directly binds mGluR1 and is preferentially enriched in large spine heads (Kalinowska et al, 2015). We reasoned that Synpo might be physically tethered to mGluR1 *via* a multivalent complex contributing to its stabilization in spines. Co-immunoprecipitation assays show that anti-mGluR1 pulls down Synpo from brain cortex extracts of adult WT mice (**Fig. 6*A***) whereas by immunolabeling mGluR1 can be detected in close apposition with Synpo in hippocampal neurons (**Fig. 6*B***).

**Figure 6.**
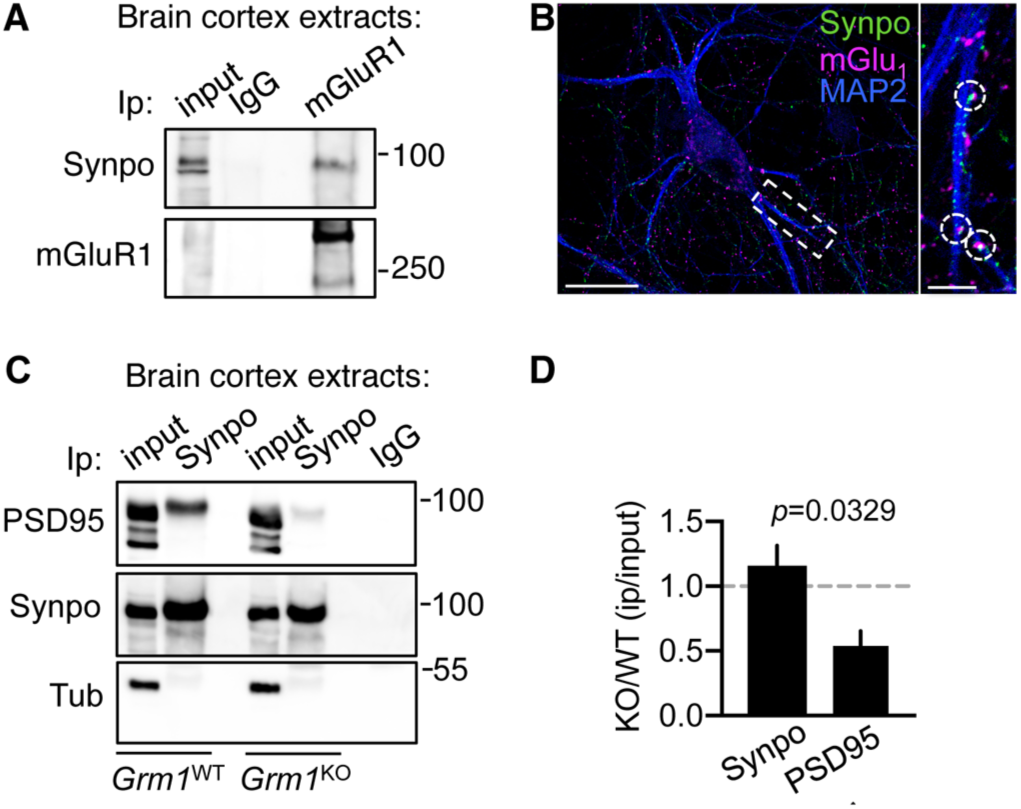
mGluR1 enables Synpo stabilization at excitatory spine synapses. A. Synpo co-precipitates with mGluR1: representative immunoblot probed for Synpo of immunoprecipitation (Ip) with anti-mGluR1 or control IgG from lysates of brain cortex of adult WT mice. B. Representative images of hippocampal neurons labeled for mGluR1, Synpo and MAP2: mGluR1 puncta are found in apposition to Synpo. Scale bar 50 µm; inset, 10 µm. C. Co-precipitation of PSD95 with Synpo in *Grm1*^WT^ and *Grm1*^KO^ mice: representative blot probed for PSD95 of immunoprecipitation with anti-Synpo antibody from lysates of brain cortex of adult *Grm1*^WT^ and *Grm1*^KO^ mice. Input, 5% of Ip lysate; Tubulin (Tub), loading control. D. Quantification of PSD95 co-precipitation with Synpo from blots as in (b). Mean ± SE, KO/WT PSD95 0.54 ± 0.12, KO/WT Synpo 1.2 ± 0.16, N=3 mice per group, two-tailed unpaired t-test.

Synpo is retrieved in the postsynaptic density (PSD) fraction of excitatory synapses (Bayés et al, 2012) and was shown to be linked to core scaffold proteins associated with the PSD (Li et al, 2017). To test whether mGluR1 contributes to Synpo stabilization at excitatory spine synapses, we used mGluR1 knockout mice (*Grm1*^KO^) to probe Synpo association with the PSD in absence of the receptor. Immunoprecipitation of Synpo from brain cortex of adult wild type littermates (*Grm1*^WT^) effectively retrieved PSD95, a core component of the PSD (**Fig. 6*C***): in contrast, pull-down of PSD95 with Synpo were decreased in *Grm1*^KO^ mice (**Fig. 6*C,D***) in absence of significant alterations in the relative abundance of the proteins (mean ± SE, *Grm1*^WT^ Synpo 2.2 ± 0.25, *Grm1*^KO^ 1.7 ± 0.21 N=3 mice per group, *p* = 0.1897; *Grm1*^WT^ PSD95 2.5 ± 0.64, *Grm1*^KO^ 2.7 ± 0.82 N=3, *p* = 0.8641, two-tailed unpaired t-test), suggesting defects in the recruitment or retention of Synpo to spine synapses.

To examine if impaired association of Synpo with the PSD may be linked to spine abnormalities in *Grm1*^KO^ mice, we used DiIC_18_ labeling to visualize dendritic spines in brain cortex tissue sections from juvenile (∼1.5-month old) and adult (7 to 12-month old) *Gmr1*^KO^ and *Grm1*^WT^ littermates. We found that in *Grm1*^KO^ mice, the density of large mushroom spines was significantly reduced compared to WT littermates (**Fig. 7*A,B***): moreover, the heads of remaining mushroom spines in *Grm1*^KO^ mice were significantly smaller compared to *Grm1*^WT^ littermates (**Fig. 7*A,C***). Collectively, our findings indicate that, when activated during mGluR-LTD, mGluR1 promotes Synpo degradation in dendrites whereas it locally supports Synpo stabilization in spines, possibly through formation of a multivalent protein assembly.

**Figure 7.**
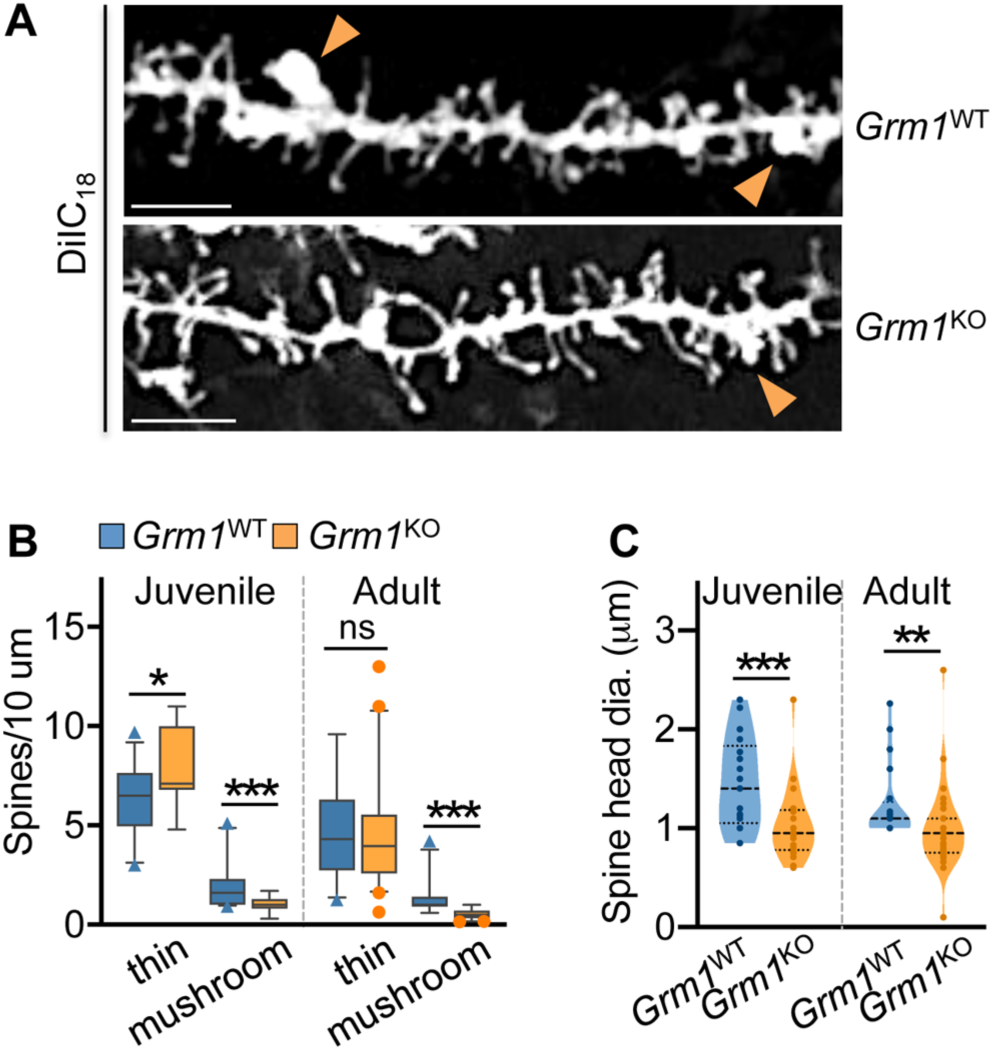
Reduced density and growth of mushroom spines in mice lacking mGluR1. A. Representative images of DiIC_18_-labeled dendritic branches in the brain cortex of adult *Grm1*^WT^ and *Grm1*^KO^ mice; scale bars 10 µm. B. Quantification of spine density per 10 µm in the cortex of *Grm1*^WT^ and *Grm1*^KO^ mice from images as in (a). Juvenile (∼1.5 month-old), mean ± SD thin spines WT 6.8 ± 1.7 (n=32 dendritic branches), KO 7.8 ± 1.9 (n=25), (*) *p*=0.025; mushroom spines WT 1.9 ± 0.98 (n=34), KO 1.10 ± 0.39 (n=19), N=2 mice per group; (***) *p*<0.001. Adult mice (∼7-12 month-old), mean ± SD thin spines WT 4.7 ± 2.2 (n=36), KO 4.4 ± 2.5 (n=46), (ns) *p*=0.29; mushroom spines WT 1.4 ± 0.87 (n=31), KO 0.54 ± 0.25 (n=39), WT N=5, KO N=4 mice (***) *p*<0.001, Mann-Whitney test. C. Quantification of mushroom spine head dimensions (diameter) in *Grm1*^WT^ *Grm1*^KO^ mice. Juvenile, mean ± SD, WT 1.5 ± 1.7µm (n=21), KO 1.0 ± 0.36µm (n=25) (***) *p*=0.001; adult, mean ±SD, WT 1.2 ± 0.38 (n=20), KO 1.0 ± 0.40 (n=40), (**) *p*=0.003, Mann-Whitney test.

## Discussion

We report that mGluR-LTD is compromised in mice lacking Synpo/SA. Interestingly, it was previously shown that in *Synpo*^KO^ mice basal neurotransmission is unaffected (Deller et al, 2003) and NMDAR-dependent LTD proceeds normally (Zhang et al, 2013). Unlike NMDAR-dependent LTD, mGluR-LTD requires rapid protein synthesis (Huber et al, 2000). Polyribosomes are present in proximity to the SA (Ostroff et al, 2010; Spacek & Harris, 1997) which was proposed to function as satellite secretory station based on the presence of proteins of the secretory pathway within the SA (Pierce et al, 2001). Thus spines containing Synpo/SA may enable local biosynthesis of proteins that support induction and expression of mGluR-LTD. Other forms of synaptic plasticity were shown to be impaired in *Synpo*^KO^ mice, including LTP (Deller et al, 2003; Jedlicka et al, 2009; Zhang et al, 2013) and homeostatic potentiation (Vlachos et al, 2013), congruent with observations that dendritic spines of *Synpo*^KO^ mice do not undergo structural expansion during potentiation (Korkotian et al, 2014; Vlachos et al, 2009), which in turn requires protein synthesis (Yang et al, 2008).

Here we show that in hippocampal neurons, mGluR-LTD selectively reduces the overall density of mushroom spines that lack Synpo, but not the number of Synpo-positive spines. In naïve mature neurons *in vitro*, Synpo is targeted to spines with a frequency similar to that reported *in vivo* (Verbich et al, 2016). This is in line with findings that the diversity of excitatory synapses of varying strength observed *in vivo* is recapitulated in neurons *in vitro* without prior history of activity, implicating cell-intrinsic mechanisms in the generation of spine diversity (Hazan & Ziv, 2020). Spines containing sER, and presumably Synpo, possess higher synaptic strength (Holbro et al, 2009). Moreover spines harboring Synpo undergo slower constitutive turnover and as such are more stable compared to mushroom spines in which Synpo is absent (Yap et al, 2020). By analogy, in the preparation used here, mGluR-LTD appears to promote loss of spines that are ‘weaker’ and ‘unstable’ *vs*. strong and stable Synpo-positive spines that harbor the SA.

The impact of mGluR-LTD on spine shrinkage/elimination is controversial. Inhibition of Gp1 mGluRs was shown to limit the shrinkage of large spines induced by low-frequency glutamate uncaging (Oh et al, 2013). In addition, chemical mGluR-LTD was shown to promote shrinkage and long-lasting spine elimination in hippocampal CA1 pyramidal neurons (Ramiro-Cortés & Israely, 2013). Both findings supported the view that mGluR-LTD produces changes in spine structural plasticity but a recent study failed to observe such changes during DHPG-LTD using similar experimental strategies (Thomazeau et al, 2020). Although the reasons for the discrepancy remain unclear, our observations reveal a previously unappreciated complex function of mGluR-LTD in the elimination of selected spines distinguished by their molecular composition, an event that cannot be captured by examination of morphology and/or dimensions alone.

Synpo expression is developmentally regulated and is modified by activity patterns that produce plasticity, including LTP and homeostatic synaptic scaling. Here we find that mGluR-LTD triggers Synpo degradation *via* activation of mGluR1. While the function of Gp1 mGluRs in promoting protein synthesis is extensively documented (Osterweil et al, 2010; van Gelder et al, 2020), their role in proteolysis is not fully understood. Gp1 mGluRs are known to regulate the expression of Fragile X Mental Retardation Protein (FMRP) by stimulating both its rapid proteasome-dependent degradation and synthesis (Hou et al, 2006; Nalavadi et al, 2012; Zhao et al, 2011). Furthermore, although proteasome-dependent and –independent proteolysis is critical for mGluR-LTD (Citri et al, 2009; Zhu et al, 2017), the identity of the substrates and impact of their degradation are largely unknown. We propose that mGluR-LTD shunts Synpo degradation to the lysosomal pathway that patrols dendrites and disposes of proteins aggregates. Degradation of dendritic Synpo may deplete a protein ‘reservoir’ otherwise available for recruitment to spines. sER tubules invade and retract from spines (Perez-Alvarez et al, 2020; Wagner et al, 2011) and, although untested, dendritic Synpo associated with sER could be co-recruited to spines. A potential scenario is that by reducing dendritic Synpo availability, mGluR-LTD may exert the dual function of eliminating ‘unstable’ spines and limit spurious spine strengthening by *de novo* recruitment of Synpo.

We find that mGluR1 is critical for controlling both the abundance of Synpo in dendritic shafts and Synpo maintenance at excitatory spine synapses, potentially via stabilization through formation of a multivalent macromolecular complex. In support of this view, we show evidence that in mice lacking mGluR1, Synpo recruitment/stabilization at excitatory synapses is impaired as indicated by its diminished association with the PSD. Consistent with this possibility, we provide the first report of profound spine dysmorphogenesis in *Grm1*^KO^ mice: in the mutants, we observed a significant reduction of mushroom spines - in both adult and juvenile mice - that also display smaller heads compared to WT littermates. Conceptually, the impact of mGluR1 both *in vivo* and in the *in vitro* neuronal network is reminiscent of what observed in circuits in the cerebellum at CF-PC synapses (Hashimoto & Kano, 2013) and adult dLGN (Narushima et al, 2016) in which mGluR1 promotes stabilization of strong synaptic connections and elimination of weaker inputs. Overall our results identify spines with Synaptopodin/ SA as the locus of mGluR-LTD and underscore the importance of the molecular anatomy of spines in synaptic plasticity.

## Author contributions

L.S.: designed research, performed research, analyzed data; Y.I.: performed research, analyzed data; C.DS: performed research, analyzed data; P.Y.W.: performed research, analyzed data; M.K.: performed research, analyzed data; R.A.M: designed research, analyzed data, wrote the paper; A.F.: designed research, analyzed data, wrote the paper.

## Acknowledgments

We thank members of the Francesconi and McKinney labs for comments on the manuscript. We would like to thank Francois Charron for his excellent technical assistance. Supported by NIMH R01MH108614 (A.F.), CIHR MOP 86724 (R.A.M.) and The Norman Zavalkoff Family Foundation (R.A.M). We acknowledge the assistance of the Neural Cell Engineering and Imaging Core of the Einstein Rose F. Kennedy Intellectual and Developmental Disabilities Research Center supported by NICHD U54 HD090260.

